# Identifying drug response by combining measurements of the membrane potential, the cytosolic calcium concentration, and the extracellular potential in microphysiological systems

**DOI:** 10.1101/2020.05.29.122747

**Authors:** Karoline Horgmo Jæger, Verena Charwat, Sam Wall, Kevin E. Healy, Aslak Tveito

## Abstract

Cardiomyocytes derived from human induced pluripotent stem cells (hiPSC-CMs) offer a new means to study and understand the human cardiac action potential, and can give key insight into how compounds may interact with important molecular pathways to destabilize the electrical function of the heart. Important features of the action potential can be readily measured using standard experimental techniques, such as the use of voltage sensitive dyes and fluorescent genetic reporters to estimate transmembrane potentials and cytosolic calcium concentrations. Using previously introduced computational procedures, such measurements can be used to estimate the current density of major ion channels present in hiPSC-CMs, and how compounds may alter their behavior. However, due to the limitations of optical recordings, resolving the sodium current remains difficult from these data. Here we show that if these optical measurements are complemented with observations of the extracellular potential using multi electrode arrays (MEAs), we can accurately estimate the current density of the sodium channels. This inversion of the sodium current relies on observation of the conduction velocity which turns out to be straightforwardly computed using measurements of extracellular waves across the electrodes. The combined data including the membrane potential, the cytosolic calcium concentration and the extracellular potential further opens up for the possibility of accurately estimating the effect of novel drugs applied to hiPSC-CMs.

## 1 Introduction

In recent reports [1, 2] we have demonstrated how microphysiological systems utilizing human induced pluripotent stem derived cardiomyocytes (hiPSC-CMs)[3, 4] can be used to estimate drug induced changes to the cardiac action potential using computational approaches. These methods use optical measurements of the membrane potential and the cytosolic calcium concentration to quantitate changes in underlying ion channel conductances and calcium handling pathways using a mathematical model of the of hiPSC-CMs dynamics. We have further shown how these estimates, at least in principle, carry over from immature cells to adult cardiomyocytes. This methodology provides information on a number of the major ion channels and when compared to data presented in [5, 6, 7, 8, 9, 10, 11], the method is able to provide reasonable estimates of the IC50 values of well-known drugs like Nifedipine, Lidocaine, Cisapride, Flecainide and Verapamil; see Table 3 of [2]. These drug affects the *I*_CaL_, *I*_NaL_ or *I*_Kr_ currents and the effect is well estimated by our methodology.

However, difficulties remain in the characterization of the fast sodium current, *I*_Na_. This is a major issue since this current more or less completely governs the rapid upstroke of the action potential and thus also the conduction velocity. Therefore, it is of great importance to characterize the effect of drugs on this current. The reason for this deficiency in our methodology is the time resolution of the data obtained by fluorescence; the data used in the inversions are provided with a resolution of 10 ms and this is far too coarse to be able to estimate the strength of *I*_Na_. Time resolution can be improved but at the cost of less accurate data and therefore another experimental technique is needed to pin down the channel density of and drug effects on *I*_Na_. It is well known that the extracellular potential can be measured in microphysical systems using multielectrode arrays; see e.g., [12, 13, 14, 15, 16]. In this report we will show that the extracellular data can be used to determine the sodium current. And therefore, by combining imaging data for the membrane potential (V) and the cytosolic calcium concentration (Ca) with data for the extracellular potential (U), we are able to identify both the fast sodium current and other major currents characterizing the action potential of the hiPSC-CMs.

The main challenge in combining V, Ca and U data is that a spatial problem needs to be resolved. When the data are given by V and Ca only, we have simply used a data trace obtained by taking the average over the whole chip (see [3, 1]) and the inversion of the data has amounted to estimating parameters describing a system of ordinary differential equations. But when U is added to the data, the extracellular potential needs to be calculated. In our present implementation, we use the bidomain model (see e.g. [17]) for this purpose. The bidomain model has already been used for inversion of U data (but not V and Ca) by several authors; see [18, 16, 19, 20, 14]. However, it is demonstrated in [18] that the bidomain model does not provide an extracellular repolarization wave. Such a wave is clearly present in the experimental data and inhomogeneities have to be introduced in the bidomain model in order to enforce a repolarization wave. These inhomogeneities are difficult to obtain from measurements, and therefore we choose to use U only to estimate the currents involved in generating the upstroke and not the whole action potential. This turns out to determine *I*_Na_ accurately, - at least in data generated by simulations.

## 2 Methods

In this section, we describe the methods applied to identify drug response by combining measurements of the membrane potential, the cytosolic calcium concentration and the extracellular potential in microphysiological systems of hiPSC-CMs.

### 2.1 Bidomain-base model simulations

In order to represent the electrical properties of a microphysiological system, we conduct simulations of the bidomain model of the form

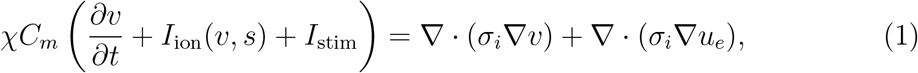

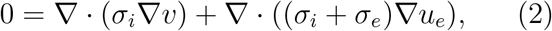

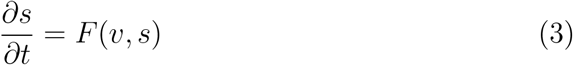

(see e.g., [21, 17, 22, 23]). Here, *υ* and *u*_*e*_ are the membrane potential and the extracellular potential, respectively. In addition, *σ*_*i*_ and *σ*_*e*_ are the bidomain conductivities of the extracellular and intracellular spaces, respectively, *C*_*m*_ is the specific membrane capacitance, and *χ* is the surface-to-volume ratio of the cell membrane. The values chosen for these parameters are given in Table 1.

**Table 1:**
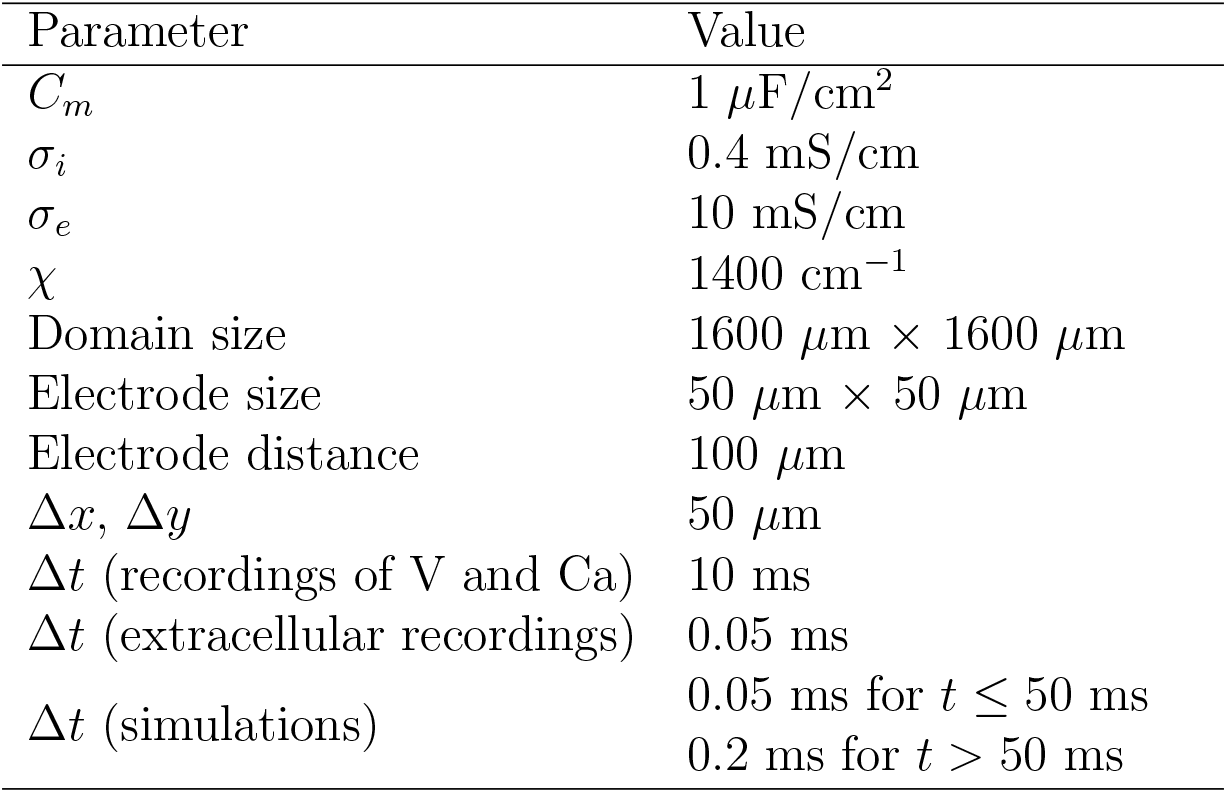
Parameter values used in the bidomain-base model simulations.

Furthermore, *I*_ion_ represents the density of currents through different types of ion channels, pumps and exchangers on the cell membrane. We use an adjusted version of the hiPSC-CM base model introduced in [2] to represent these currents. The current density is then given on the form

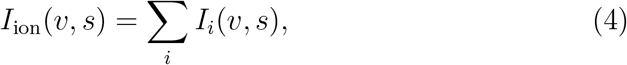

where each *I*_*i*_ represents the current through a specific type of ion channel, pump or exchanger. The hiPSC-CM base model includes a number of additional state variables representing the gating of ion channels and intracellular Ca^2+^ concentrations. These variables are represented by *s* in the model above, and their dynamics are modeled by a set of ordinary differential equations (ODEs) given by *F* (*v, s*). A number of parameters in the hiPSC-CM base model have been adjusted to make the size of eight currents of particular interest (*I*_Kr_, *I*_NaL_, *I*_CaL_, *I*_Na_, *I*_to_, *I*_Ks_, *I*_K1_, and *I*_f_) close to the size of the currents in the model described in [24], which is fitted to recordings of hiPSC-CMs from several different studies. The adjusted parameter values of the hiPSC-CM base model are given in Table 2, and the base model is, for completeness, given in the supplementary information.

**Table 2:** Parameter values used in the hiPSC-CM base model. The remaining parameter values are the same as specified in [2].

| Parameter | Value | Parameter | Value |
| --- | --- | --- | --- |
| $[\text{Ca}^{2+}]_e$ | 0.42 mM | $\bar{I}_{\text{NaK}}$ | 1.9 $\mu\text{A}/\mu\text{F}$ |
| $[\text{K}^+]_e$ | 5 mM | $\bar{I}_{\text{NaCa}}$ | 9.4 $\mu\text{A}/\mu\text{F}$ |
| $[\text{Na}^+]_e$ | 140 mM | $\bar{I}_{\text{pCa}}$ | 0.12 $\mu\text{A}/\mu\text{F}$ |
| $g_{\text{Na}}$ | 2.6 mS/$\mu\text{F}$ | $\bar{J}_{\text{SERCA}}$ | 0.00016 mM/ms |
| $g_{\text{NaL}}$ | 0.03 mS/$\mu\text{F}$ | $\alpha_{\text{RyR}}$ | 0.0052 ms<sup>-1</sup> |
| $g_{\text{to}}$ | 0.21 mS/$\mu\text{F}$ | $\beta_{\text{RyR}}$ | 0.0265 mM |
| $g_{\text{Kr}}$ | 0.075 mS/$\mu\text{F}$ | $\alpha_d^c$ | 0.0027 ms<sup>-1</sup> |
| $g_{\text{Ks}}$ | 0.0127 mS/$\mu\text{F}$ | $\alpha_{sl}^c$ | 0.3 ms<sup>-1</sup> |
| $g_{\text{K1}}$ | 0.05 mS/$\mu\text{F}$ | $\alpha_n^s$ | 0.0093 ms<sup>-1</sup> |
| $g_{\text{f}}$ | 0.012 mS/$\mu\text{F}$ | $B_{\text{tot}}^c$ | 0.063 mM |
| $g_{\text{bCl}}$ | 0.0001 mS/$\mu\text{F}$ | $B_{\text{tot}}^d$ | 2.7 mM |
| $g_{\text{bCa}}$ | 0.0005 mS/$\mu\text{F}$ | $B_{\text{tot}}^{sl}$ | 1.45 mM |
| $g_{\text{CaL}}$ | 1.8 nL/($\mu\text{F}$ ms) | $B_{\text{tot}}^s$ | 60 mM |

The geometry of the domain used to represent a microphysiological system is decsribed in Figure 1 and Table 1. We consider a two-dimensional domain, and initiate a travelling wave by stimulating the lower left corner. Furthermore, 8×8 electrodes are distributed in the center of the domain, and we record the extracellular potential in these electrodes. On the boundary of the domain, we apply the Dirichlet boundary condition *u*_*e*_ = 0.

**Figure 1:**
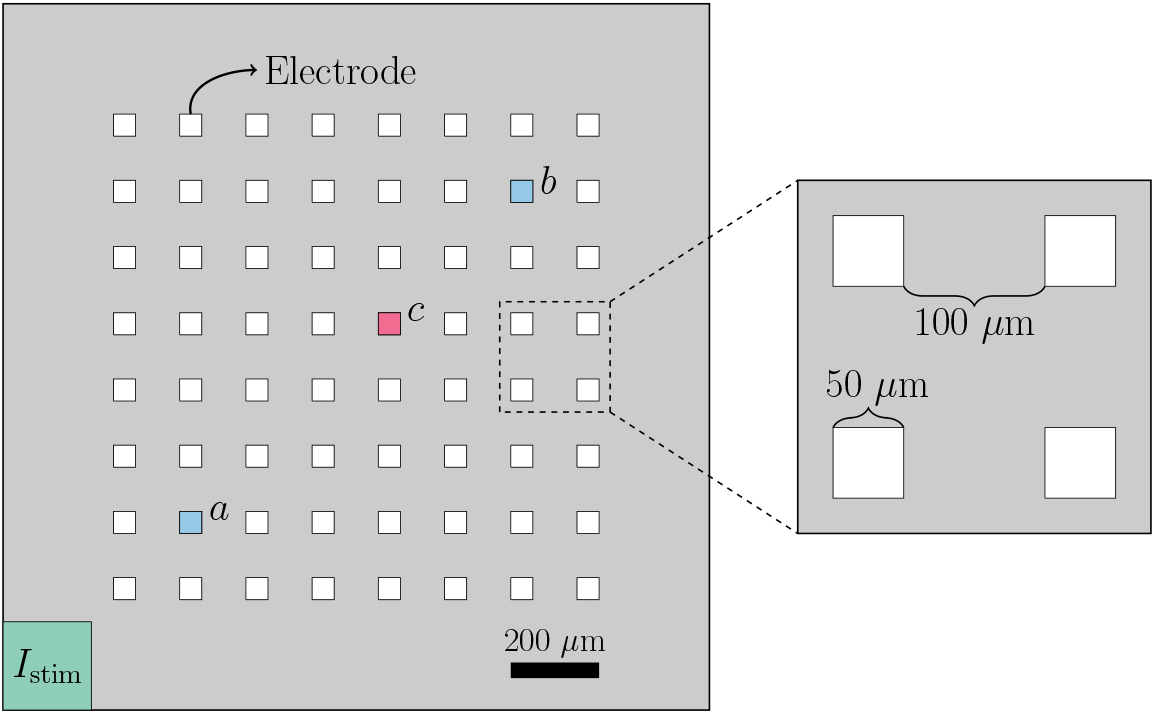
Geometry used in the bidomain-base model simulations. The domain contains 8×8 evenly spaced electrodes, and the propagating wave is initiated by stimulating the lower left corner of the domain. The conduction velocity is computed between the electrodes marked as *a* and *b*, and when we plot the extracellular potential of just a single electrode, we consider the electrode marked as *c*.

In addition to the bidomain simulations, we in some cases conduct simulations in which the spatial variation of the variables are ignored. In that case, the system (1)–(3) can be written as a pure ODE system of form

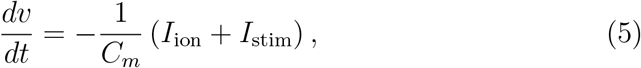

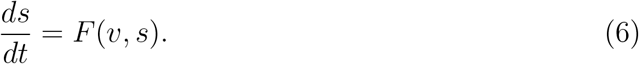

Below, we refer to the system (5)–(6) as the pure ODE system.

#### 2.1.1 Adjustment factors

In order to investigate whether we are able to identify drug responses from combined measurements of V, Ca and U data, we perform numerical simulations representing a number of different drugs reported in [7]. More specifically, we simulate ion channel blockers by introducing adjustment factors *λ*_*i*_ ≤ 0, such that *I*_ion_ is given by

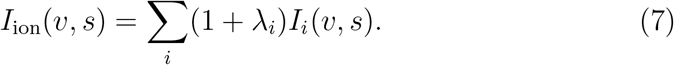

In our simulation, we consider the drug effect on three major ionic currents, *I*_Kr_, *I*_CaL_ and *I*_Na_, believed to be of importance in evaluation of drug safety (see e.g., [7]), and we therefore introduce the four adjustment factors *λ*_Kr_, *λ*_CaL_ and *λ*_Na_.

#### 2.1.2 Bidomain-base model simulations used to generate data

We wish to investigate how V, Ca and U data from a microphysiological system can be used to identify the effect of drugs. In order to generate data representing this type of recordings, we perform bidomain-base model simulations. The simulation procedure used to generate the data is illustrated in Figure 2.

**Figure 2:**
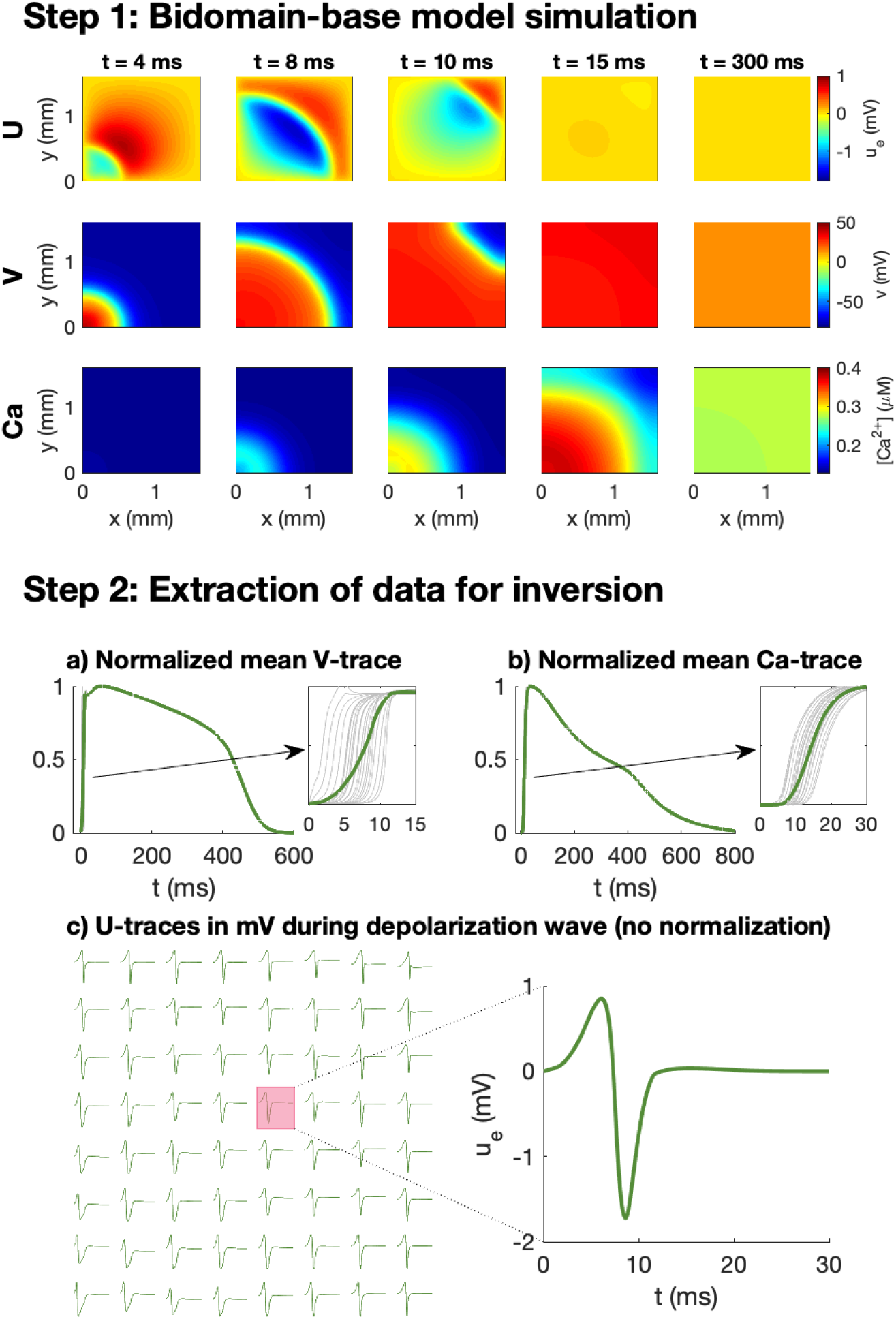
Simulation procedure used to generate data for the inversions. In Step 1, we conduct bidomain-base model simulations, recording V, Ca and U for a number of time steps. In Step 2, we extract V and Ca data for inversion by recording the mean V and Ca over the domain at each time step and normalizing the traces to values between 0 and 1. In addition, U is recorded (in mV) in 8×8 electrodes at each time step during the depolarization wave.

As illustrated in the upper panel of Figure 2, for a given combination of *λ*-values, we first perform bidomain-base model simulations for the duration of an entire action potential and store the extracellular potential, the membrane potential, and the cytosolic calcium concentration in each mesh point for each time step. We then wish to convert these recorded solutions into corresponding measurements that may be performed in microphysiological systems.

The first considered type of measurement is optical measurements of voltage-sensitive dyes. This type of measurement is mimicked in the computations by extracting the mean membrane potential over the entire domain for each time step. Because the exact conversion factor between the pixel intensity of the optical measurements and the associated membrane potential (in mV) is not known for this type of optical measurement, we normalize the mean V-trace to values between 0 and 1. Moreover, because the time resolution of the measurement equipment for the optical measurements typically is quite limited (see e.g., [1]), we only store the membrane potential at relatively large time steps (every 10 ms). Traces representing optical measurements of the cytosolic calcium concentration is generated in exactly the same manner.

In addition to the optical measurements of V and Ca, the extracellular potential may also be measured in electrodes located in the microphysiological system. In the simulations, we extract this type of measurements by storing the mean extracellular potential in the grid points overlapping the location of the electrodes. We are primarily interested in the extracellular potential during the depolarization wave, and we therefore only store the extracellular potential in the beginning of the simulation (typically the first 50 ms).

#### 2.1.3 Pure ODE simplification used in the inversion procedure

During the inversion procedure, we wish to identify the *λ*-values associated with some considered data, generated as explained in the previous section. In this inversion procedure, we need to conduct simulations using a large number of different *λ*-values for comparison to the data. As mentioned above, the repolarization wave is known to be poorly represented by the bidomain model when applied to small collections of hiPSC-CMs. Therefore, we perform spatial simulations only for the start of the action potential. The data from this part of the simulation is used to determine the conduction velocity, the Ca transient time to peak and the terms of the cost function related to the extracellular potential, U. By performing spatial simulations only for the beginning of the action potential, we save time by avoiding a full bidomain simulation for all of the different parameter combinations. To generate the full V and Ca traces, we instead run a simple pure ODE simulations of the form (5)–(6) of the full action potential. The solution of this ODE simulation is used to compute all the terms in the cost function (see Section 2.2.1), except for the ones related to U, the Ca transient time to peak and the conduction velocity.

#### 2.1.4 Stimulation protocol and update of initial conditions

In the pure ODE simulations, the cell is stimulated by a constant stimulus current of −5 *μ*A/cm^2^ until *υ* is above −40 mV. In the bidomain simulations, we stimulate the domain in the lower left corner as illustrated in Figure 1 by a 5-ms-long constant membrane stimulus current of −40 *μ*A/cm^2^. Note that after each parameter change, and before any bidomain or pure ODE simulation is performed, we conduct an ODE simulation for 10 AP cycles, stimulating at 1 Hz, to update the initial conditions.

#### 2.1.5 Numerical methods

The numerical simulations of the bidomain model are performed using an operator splitting procedure (see e.g., [25, 23, 26]). For each time step, we first solve the non-linear part of the system (i.e., (1) and (3) with the left-hand side of (1) set to zero), using the GRL2 method [27]. Then, we solve the remaining linear part of the system (i.e., (1) and (2) with *I*_ion_ = 0) using a finite difference discretization in space and a backward Euler discretization in time. The discretization parameters used in the numerical simulations are given in Table 1. In the pure ODE simulations used in the inversion procedure we apply the same GRL2 method as in the non-linear part of the bidomain simulations, and in the pure ODE simulations used to update the initial conditions after a parameter change, we use the *ode15s* ODE-solver in Matlab.

### 2.2 Inversion procedure

In this section, we will describe the inversion procedure applied to identify the effect of drugs based on V, Ca and U data in the form of adjustment factors, *λ* (see Section 2.1.1).

#### 2.2.1 Cost function

In the inversion procedure, we wish to find appropriate adjustment factors, *λ*, such that the solution of the model specified by *λ* is as similar as possible to the considered data. To this end, we define a cost function *H*(*λ*), measuring the difference between the data and the model solution. This cost function is defined as

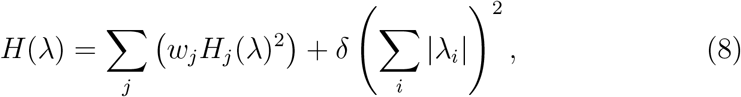

where each *H*_*j*_(*λ*) represent various differences between the data and the model solution specified by *λ*, and *w*_*j*_ are weights for each of these terms. In addition, *δ* is the weight of the regularization term (Σ_*i*_ |*λ*_*i*_|)^2^, which is included so that small adjustments, *λ*, are preferred if several choices of *λ* result in almost identical solutions. The individual cost function terms, *H*_*j*_(*λ*), are defined below.

##### Action potential and calcium transient durations

A number of terms in the cost function measure differences in the action potential and calcium transient durations. These terms take the form

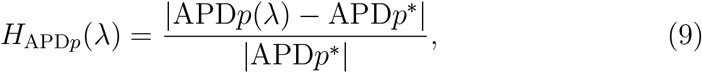

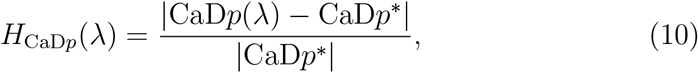

for *p* = 20, 30, …70, 80 The APD*p* value is measured as the time from V is *p*% below its maximum value during the upstroke of the action potential to the time at which V reaches a value below *p*% of the maximum during the repolarization phase; see Figure 3. APD*p*(*λ*) is the value obtained from the solution of the model given by the parameter vector *λ*, whereas APD*p*^∗^ is the value obtained from the data. The calcium transient durations, CaD*p*, are defined just like the action potential durations. Note also that the notation of a ∗ marking the data values is used for all the terms in the cost function (see below).

**Figure 3:**
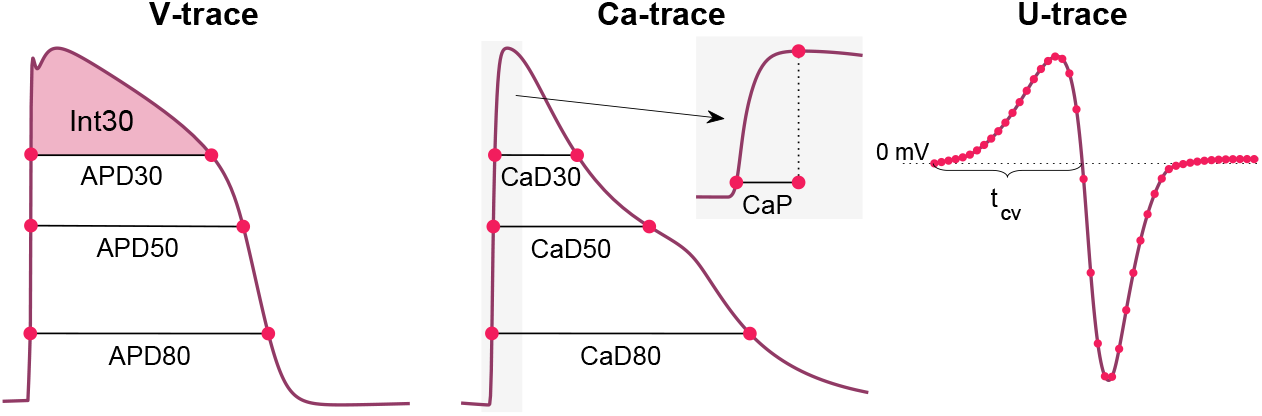
Illustration of some of the properties included in the cost function (8). From the V-trace, we include a number of APD-values (see (9)), and the integral of V above APD30 (see (11)). From the Ca-trace, we include a number of CaD-values (see (10)) in addition to the Ca transient time to peak, CaP (see (13)). From the U data, we include the value in each time point and each electrode (see (14)) and the conduction velocity computed using the time points in which U crosses 0 mV after the peak in different electrodes (see (15) and Figure 1).

##### Membrane potential integral

The cost function also includes a term of the form

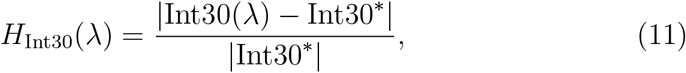

where Int30 is defined as

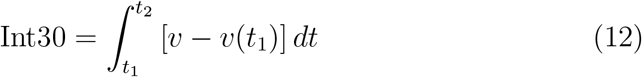

and *υ* is the membrane potential. The values *t*_1_ and *t*_2_ are here time points corresponding to when V is 30% below the maximum value during the depolarization and repolarization phases of the action potential (see Figure 3).

##### Calcium transient time to peak

The typical time resolution of the V and Ca data (10 ms) is too coarse to be able to detect changes in the upstroke velocity of the action potential. However, the upstroke of the calcium transient is slower, and therefore, changes in the upstroke of the calcium transient may be detected. To measure this upstroke velocity, we include a term of the form

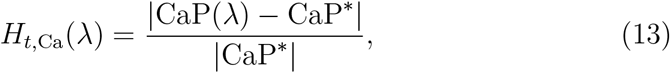

where CaP is the time from Ca is 10% above its minimum value until it reaches its maximum (see Figure 3).

As mentioned above, we use a bidomain simulation instead of a pure ODE simulation to compute an estimate of CaP(*λ*) (and CaP^∗^ in the case of simulated data). We conduct this bidomain simulation for a rather short time interval (typically 50 ms) and record Ca in all the grid points. Then, we extract the grid points that have reached their peak Ca concentration during this simulation, and set up a normalized mean Ca-trace for these grid points. The value of CaP is then computed from this normalized mean Ca-trace. When the data is generated, we use the same procedure of considering the solution for only the first part of the simulation (e.g., 50 ms) when we compute CaP^∗^.

##### Extracellular potential

In order to detect changes in U measured at the electrodes, we include a cost function term of the form

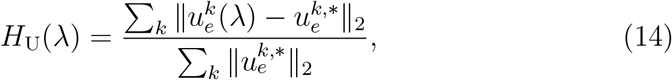

where 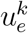 is a vector denoting U (in mV) in electrode *k* for each of the recorded time steps.

##### Conduction velocity

In addition, the cost function includes a term for the conduction velocity of the form

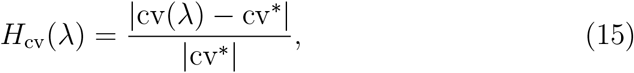

where cv denotes the conduction velocity (in cm/s) computed from U recorded in the electrodes. More specifically, the conduction velocity is computed as the distance between the center of the two electrodes marked as *a* and *b* in Figure 1 divided by the time between U crosses 0 mV after the peak in these two electrodes (see Figure 3).

##### Cost function weights

In the applications of the inversion procedure reported below, we have used the weight *w*_*j*_ = 1 for each of the cost function terms, except for *H*_APD80_ and *H*_CaD80_, which are given the weight of 5. In addition, we use the value *δ* = 0.01 for the regularization term. All terms of the cost function are scaled and thus have no unit, and therefore also *w* and *δ* are unit free.

#### 2.2.2 Continuation-based optimization

In order to find the optimal *λ*-values minimizing the cost function *H* defined in (8), we apply a continuation-based minimization procedure. This approach is described in detail in [2]. In short, we attempt to find the optimal *λ*-values by gradually moving from a known solution *λ*^0^ = 0 to the final *λ* fitting the data as best possible. To this end, we introduce a parameter *θ* that is gradually increased from 0 to 1, and define an intermediate cost function

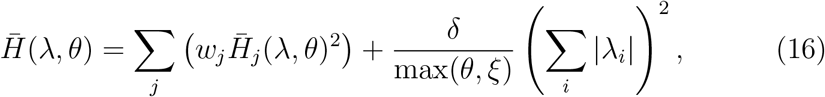

where 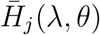 are adjusted versions of each of the cost function terms defined above and *ξ* is some small number (e.g., 10^−10^). In general, 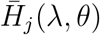 is defined as

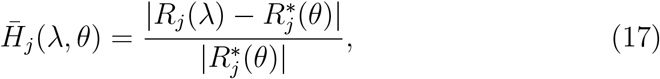

where *R*_*j*_ is some property of the solution, for example an action potential duration. Moreover, *R*_*j*_(*λ*) is the property computed in the solution of the model defined by *λ*, and 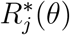 is defined as

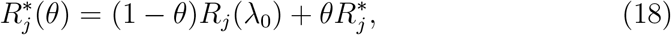

where *R*_*j*_(*λ*_0_) is the property computed from the model solution defined by *λ*^0^ = 0, and 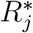 is the property computed from the data we are trying to invert.

Considering the cost function terms, *H*_*j*_, defined above, all the terms may straightforwardly be defined on the form (17) except for the term *H*_U_ defined in (14). For this term, we instead define

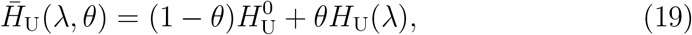

where 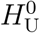 is defined by (14) with 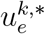 replaced by the model solution 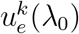, and *H*_U_(*λ*) is defined by (14).

From the definitions (16)–(19), we observe that for *θ* = 0, 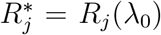 and 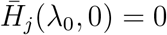, implying that the known solution *λ* = *λ*_0_ minimize (16) for *θ* = 0. Furthermore, for *θ* = 1, 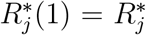 defined by the data, 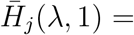 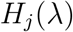, and 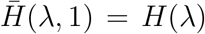. Consequently, for *θ* = 1, the minimum of 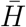 defined in (16) is the same as the minimum of *H* defined in (8).

The advantage of the continuation algorithm is that we can move from the known optimal value *λ*_0_ to the final optimal value *λ* gradually in a number of iterations, and that we for each iteration can assume that the new estimated *λ* is in the vicinity of the optimal *λ* from the previous iteration. In the applications of the inversion procedure reported below, we use four *θ*-iterations in the continuation algorithm (*θ* = 1/4, *θ* = 1/2, *θ* = 3/4 and *θ* = 1). In the first three iterations, *m*, we draw 63 random initial guesses of *λ*(*θ*_*m*_) in the vicinity of *λ*(*θ*_*m*−1_), and in the last iteration, we draw 126 initial guesses. More specifically, the initial guesses for *λ*_*i*_(*θ_m_*) are drawn from [*λ*_*i*_(*θ*_*m*−1_) − 0.2, *λ*_*i*_(*θ*_*m*−1_) + 0.2], where *λ*_*i*_(*θ*_0_) is defined to be 0. From these initial guesses we minimize the cost function 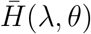 using the Nelder-Mead algorithm [28]. In the first three *θ*-iterations, we use 10 iterations of the Nelder-Mead algorithm for each initial guess, and in the last *θ*-iteration, we use 25 iterations.

### 2.3 Adjustment of extracellular concentrations

In order to better distinguish between different ion channel blockers, we consider the drug effects under different extracellular conditions. In particular, we vary the extracellular calcium concentration by introducing a number of known adjustment factors *γ*_Ca_ such that

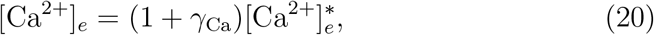

where [Ca^2+^]_*e*_ is the extracellular calcium concentration and 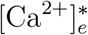 is the default extracellular calcium concentration reported in Table 2.

In the inversions reported below, we use the two values *γ*_Ca,1_ = 0 and *γ*_Ca,2_ = 0.25. We generate V, Ca and U data using the approach described in Section 2.1.2 for both of these values of *γ*_Ca_. Furthermore, for a given choice of *λ* in the inversion procedure, we compute a version of the solutions for each of these *γ*_Ca_-values, and define the cost function

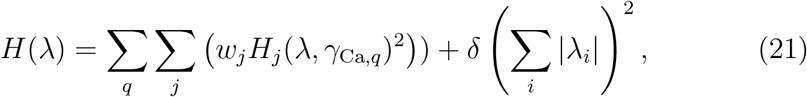

where *q* counts the different extracellular conditions and *H*_*j*_ (*λ, γ*_Ca*,q*_) represents the cost function terms computed using the extracellular calcium concentration defined by *γ*_Ca*,q*_.

### 2.4 Measuring the extracellular potential

Experimentally, the extracellular potential was measured in a monolayer of cardiomyocytes using the microelectrode array system MED64-Basic with P515 electrode dishes. Cardiomyocytes (CM) were differentiated from the human induced-pluripotent stem cell (hiPSC) line Wild Type C (WTC, Coriell Repository, # GM25256) expressing the fluorescent calcium reporter GCaMP6f [29]. Cells were differentiated into CM using a modified version of the Palecek protocol [30] applying 6 *μ*m CHIR, 5 *μ*M IWP4 and minus insulin media for 48 h each. MED64 electrode dishes were coated with polyethyleneimine followed by Matrigel and handled as suggested in the vendor application notes [31, 32]. CM were seeded to form a confluent cell layer and allowed to stabilize for 30 days with media changes of RPMI 1640 + B27 supplement (Gibco) every 2-3 days. Flecainide (Abcam, ab120504) doses were prepared in RPMI 1640 + B27 from a 25 mM stock in DMSO. Each drug dose (0, 1, 2.5 and 10 *μ*M flecainide) was incubated for at least 30 min before measurements. Extracellular potential recordings were performed using the Mobius software (Version Win 7 0.5.1) with the template for spontaneous QT recording. Traces of spontaneous beating activity were recorded on all 64 electrodes and directly exported as raw data without any pre-filtering or peak extraction.

## 3 Results

In this section, we report the results of some applications of the inversion procedure defined above. First, we investigate the sensitivity of the V, Ca and U data in response to perturbations of the *I*_Kr_, *I*_CaL_ and *I*_Na_ currents, and how this sensitivity may be increased when data for several different extracellular calcium concentrations are included. Next, we show some examples of extracellular repolarization waves, and explain why we have chosen to only consider the extracellular depolarization waves in the inversion procedure. Afterwards, we show some examples of how bidomain-base model simulations are able to reproduce measured drug induced effects on the conduction velocity, illustrating that including U data in the inversion procedure improves the identifiability of drug effects on *I*_Na_. Finally, we test the inversion procedure by using it to identify drug effects for a number of simulated drugs and investigate how the accuracy of the inversion is affected when noise is included in the simulated data.

### 3.1 Sensitivity of currents

In Figure 4, we investigate the effect on the simulated V, Ca and U data of perturbing the *I*_Kr_, *I*_CaL_ and *I*_Na_ currents. We compute the data for the default base model (*λ* = 0) and for two perturbations *λ*_*i*_ = −0.2 and *λ*_*i*_ = −0.4 for each of the currents *i* = Kr, CaL, Na, corresponding to 20% and 40% block of the currents, respectively (see Section 2.1.1). Note that in the upper panel (investigating block of *I*_Kr_), we have used a pacing frequency of 0.5 Hz instead of 1 Hz in order to allow for increased action potential durations resulting from block of *I*_Kr_.

**Figure 4:**
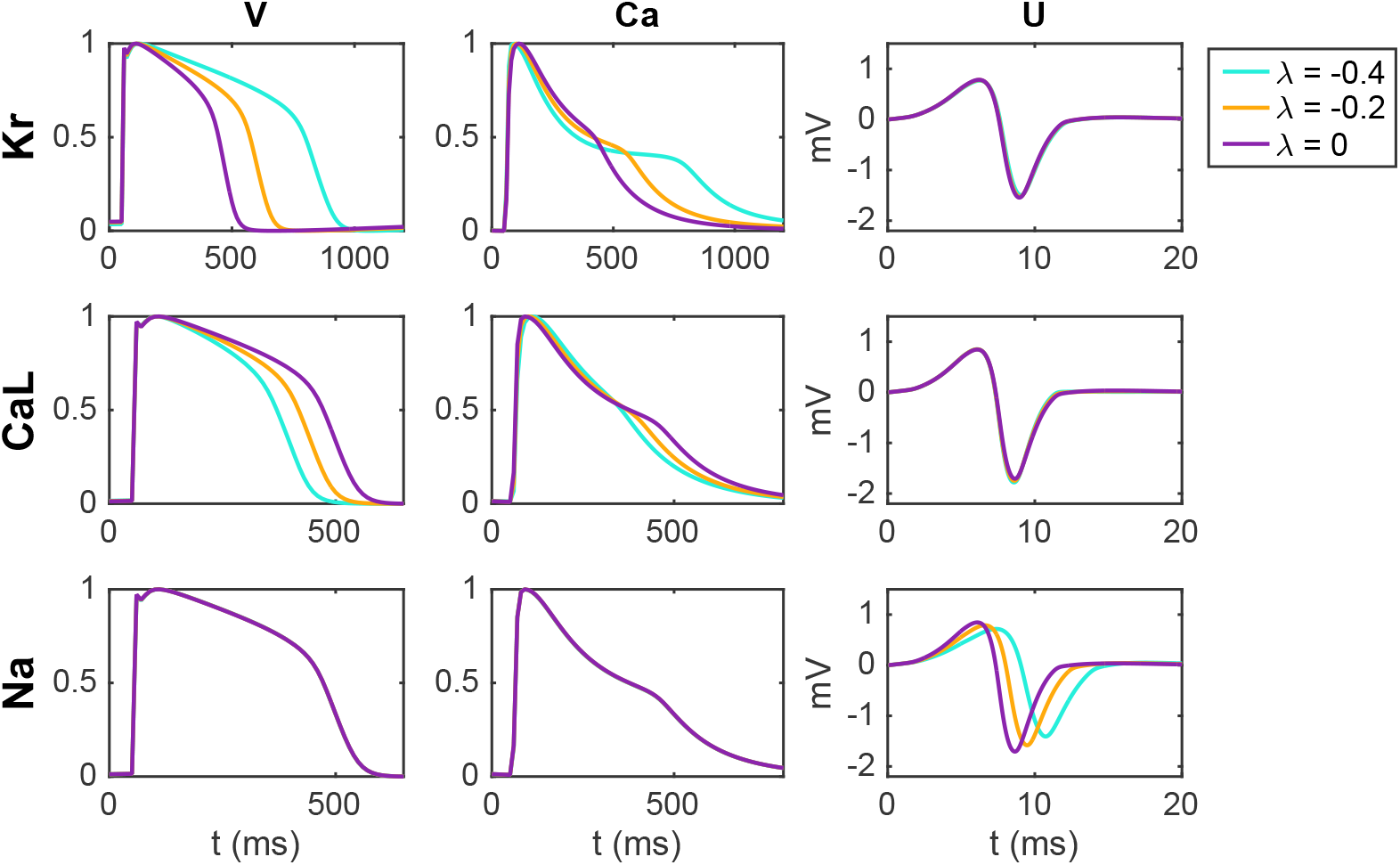
Effect of perturbing *I*_Kr_, *I*_CaL_ and *I*_Na_ on the V, Ca and U data. For the U data, we show the extracellular potential in the electrode marked as *c* in Figure 1 and the time scale is zoomed in on the first 20 ms of the simulation.

We observe that block of the *I*_Kr_ current results in increased action potential and calcium transient durations, whereas block of *I*_CaL_ results in decreased action potential and calcium transient durations. Moreover, no effects are visible in the U data resulting from block of *I*_Kr_ or *I*_CaL_ (recall that the U data is only considered for the first 20 ms). For block of the *I*_Na_ current, on the other hand, no effects on the action potential and calcium transient durations are visible, but there are clearly visible effects on the U data. Specifically, the amplitude of U is decreased and the timing of the peaks is delayed in response to block of the *I*_Na_ current.

Furthermore, in Figure 5, we report the conduction velocities computed from the U data in response to perturbations of the three currents. We observe that for perturbations of *I*_Kr_ and *I*_CaL_, the effects on the conduction velocity are quite small, whereas block of *I*_Na_ results in a considerably decreased conduction velocity.

**Figure 5:**
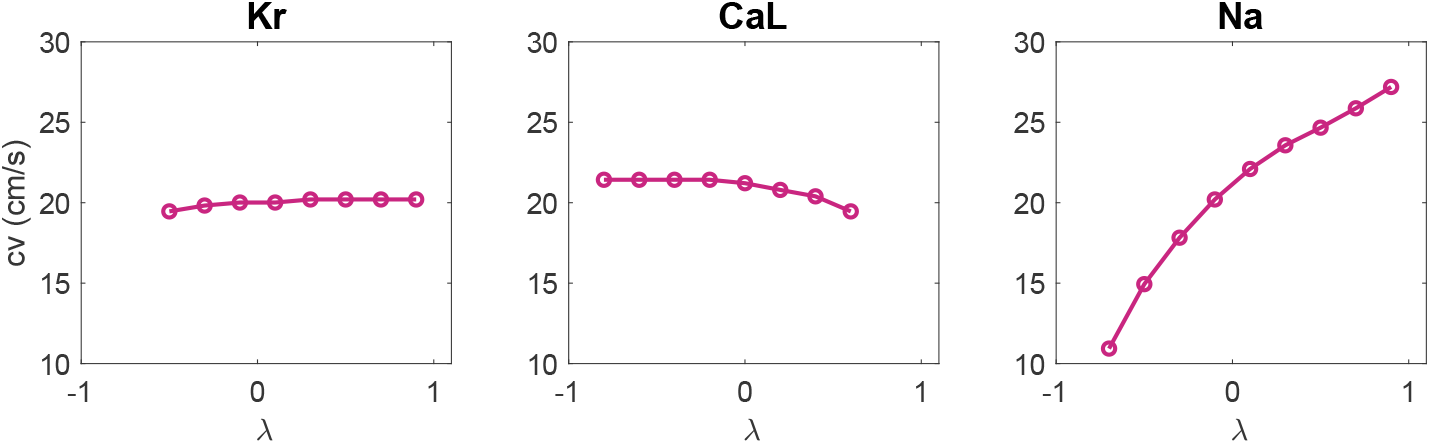
Effect of perturbing *I*_Kr_, *I*_CaL_ and *I*_Na_ on the computed conduction velocity. The conduction velocities are computed from the U data as explained in Section 2.2.1.

### 3.2 Adjustment of extracellular concentrations

Because the effect on the V, Ca and U data of blocking *I*_Kr_ is almost exactly the opposite of the effect of blocking *I*_CaL_ (see Figure 4), we expect that determining the correct combination of block for these two currents will be difficult. Therefore, it could be useful to consider drug effects under different extracellular conditions in order to increase the chance of identifying the correct block for the two currents.

Figure 6 shows the effect of block of the *I*_Kr_, *I*_CaL_ and *I*_Na_ currents for three different extracellular calcium concentrations. We consider the default extracellular calcium concentration specified in Table 2 (*γ*_Ca_ = 0), a 10% increased concentration (*γ*_Ca_ = 0.1), and a 25% increased concentration (*γ*_Ca_ = 0.25). The solid lines show the default solutions for each of these extracellular environments, and we observe that the action potential and calcium transient durations are increased as the extracellular calcium concentration is increased. Furthermore, the dotted lines show the solutions corresponding to 20% block of the considered currents. For all the considered U data and for the V and Ca data for block of *I*_Na_, the different extracellular calcium concentrations do not seem to have a significant effect. However, for the V and Ca data for block of *I*_Kr_ and *I*_CaL_, we observe that the effect of the block varies for the different concentrations. For example, the effect of block of *I*_Kr_ is more prominent for an increased extracellular calcium concentration.

**Figure 6:**
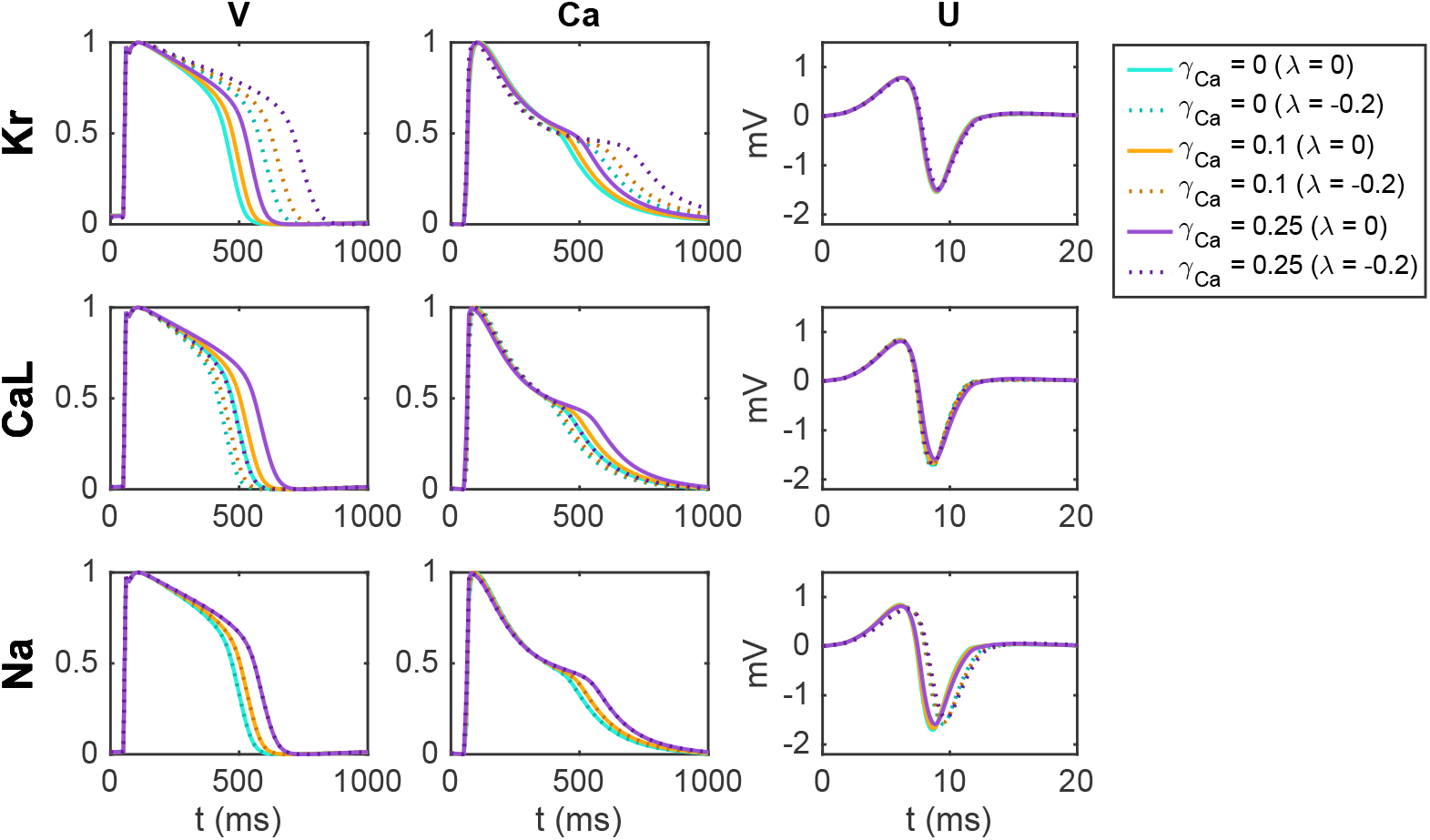
Effect of perturbing *I*_Kr_, *I*_CaL_ and *I*_Na_ on the V, Ca and U data for three different adjustments, *γ*_Ca_, of the extracellular calcium concentration (see Section 2.3). For the U data, we show the extracellular potential in the electrode marked as *c* in Figure 1 and the time scale is zoomed in on the first 20 ms of the simulation.

In Figure 7, we show an example of a case where including an additional extracellular calcium concentration could help identify the correct block of *I*_Kr_ and *I*_CaL_. In the upper panel, we use the default extracellular calcium concentration of Table 2. The solid line shows V and Ca for the default base model, and the dotted line shows the solution for a case with 20% block of *I*_Kr_ and 33% block of *I*_CaL_. We observe that V and Ca look very similar in these two cases. As a result, the inversion procedure might mistake the case of (*λ*_Kr_ = −0.2, *λ*_CaL_ = −0.33) to the case with no block. In the lower panel, however, we compare the two solutions for the case with an increased extracellular calcium concentration, and in this case, the difference between the solutions is more prominent. Consequently, the inversion procedure can more easily distinguish between the two cases when the solutions for both extracellular calcium concentrations are included.

**Figure 7:**
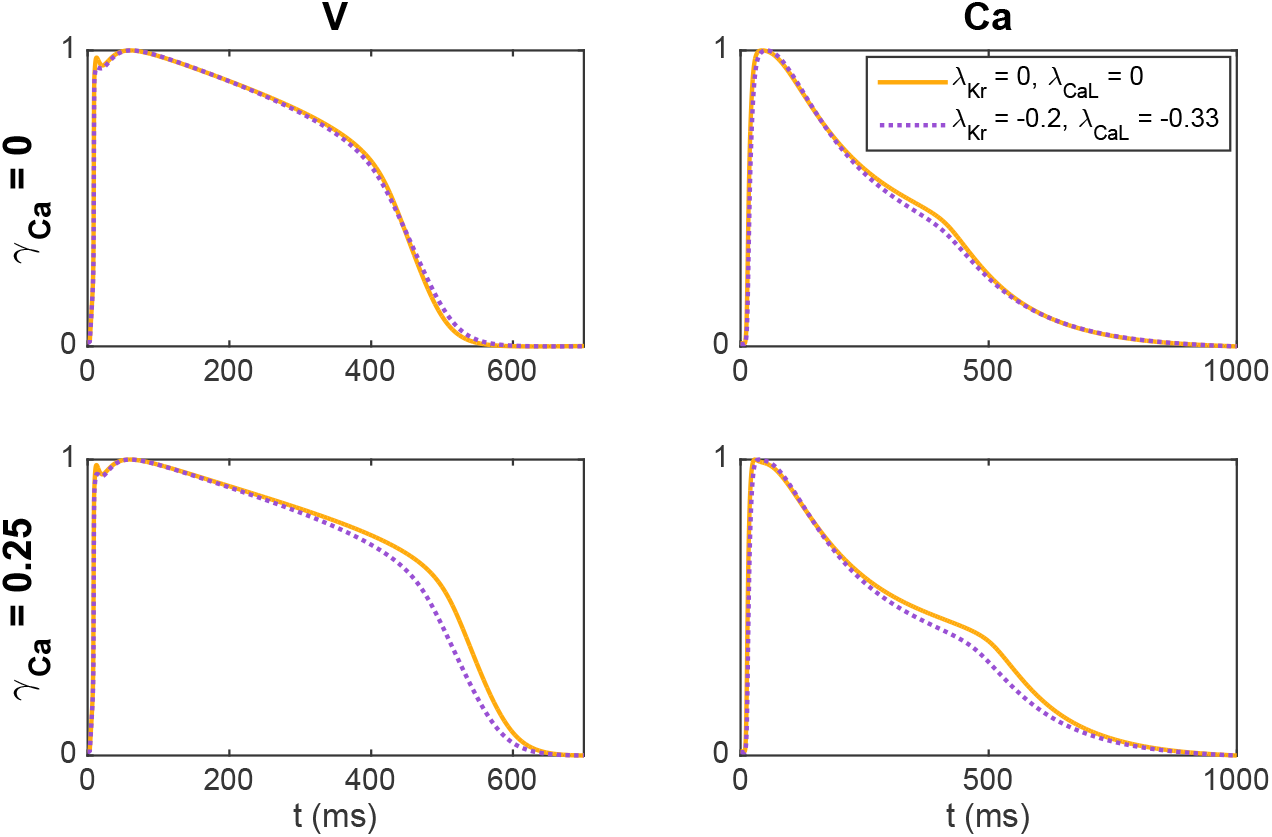
Illustration of how including an extra extracellular calcium concentration adjustment may improve the identifiability of *I*_Kr_ and *I*_CaL_. In the upper panel, we compare V and Ca for two different block combinations for the default extracellular calcium concentration (*γ*_Ca_ = 0), and in the lower panel we consider the solutions for a 25% increased extracellular calcium concentration (*γ*_Ca_ = 0.25). For *γ*_Ca_ = 0, the two solutions are very similar, but the difference is increased for *γ*_Ca_ = 0.25.

### 3.3 Estimating the action potential duration

Figure 8A shows measurements of the extracellular potential in 64 electrodes recorded for collections of hiPSC-CMs. In the upper panel, we observe that there is some early activity, corresponding to the time of the upstroke of the action potential, in addition to a weaker signal after some hundred milliseconds. This weaker signal occurs at the time of repolarization of the action potential and is therefore referred to as the repolarization wave. In the lower panel, we zoom in on this repolarization wave, and we observe that the extracellular potential reaches a magnitude of up to about 0.1-0.2 mV in this period.

**Figure 8:**
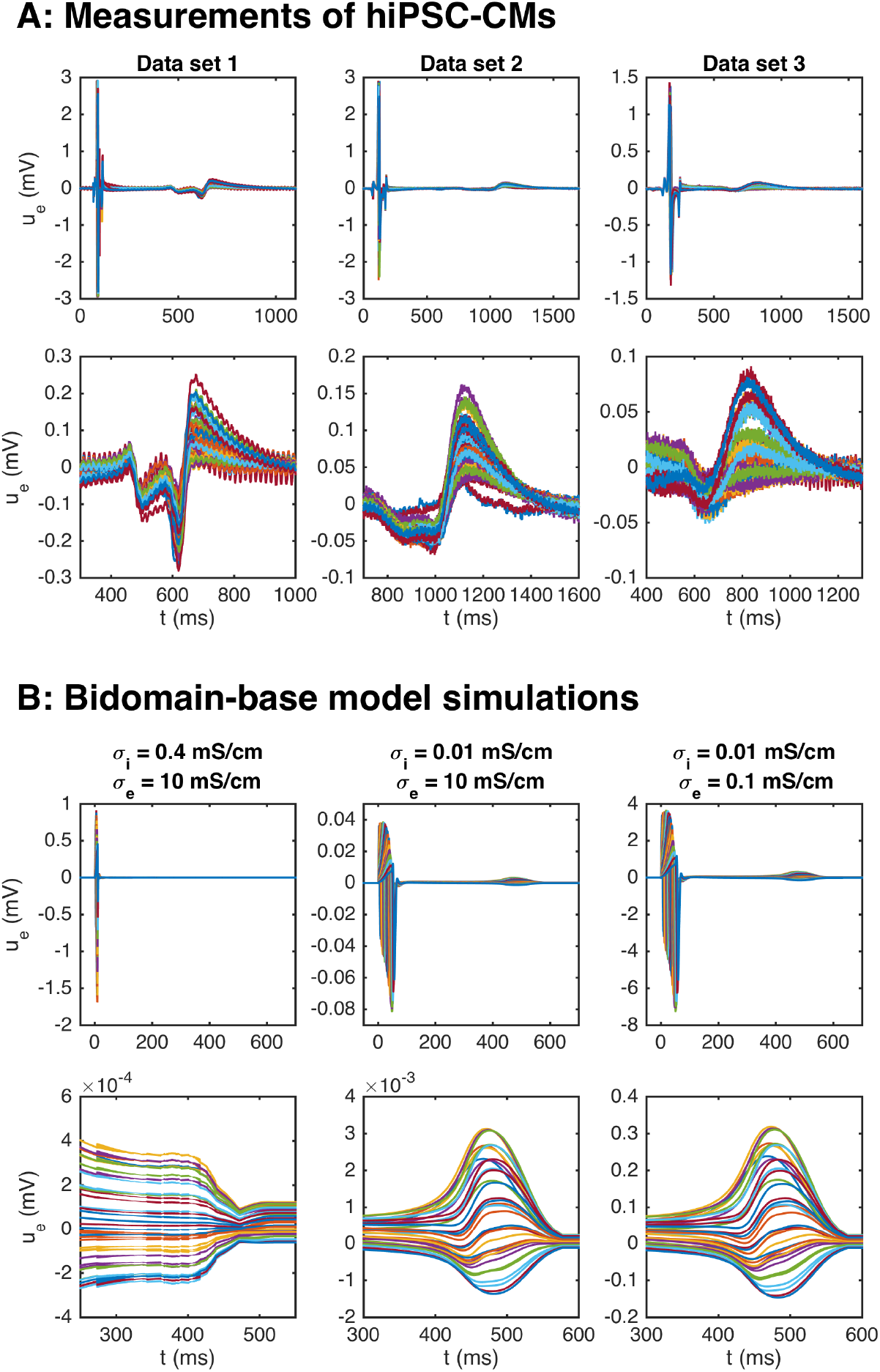
Repolarization wave in hiPSC-CM data and bidomain-base model simulations. The upper panel shows measurements of U during an action potential for three different data sets. The traces for the 64 electrodes are overlaid in the plots. In the second panel, we zoom in on the repolarization wave. The third panel shows U in three bidomain-base model simulations, and the bottom panel shows the corresponding repolarization waves. The parameters used in the simulations are given in Table 1, except for *σ*_*i*_ and *σ*_*e*_ which are reported in the plot titles.

It has been demonstrated (see e.g., [18]), that bidomain simulations of small collections of hiPSC-CMs tend to give rise to very weak or even non-exsisting repolarization waves. Indeed, in the leftmost panel of Figure 8B, we plot the extracellular potential in a bidomain-base model simulation using the default parameter values of Table 1, and we observe that the magnitude of the wave in this case is very small, and completely invisible in the upper plot over the entire action potential duration. If we decrease the intracellular conductivity, *σ*_*i*_, on the other hand, a repolarization wave is visible, but the entire extracellular signal is quite weak (see the center panel of Figure 8B). However, if the extracellular conductivity, *σ*_*e*_, is decreased as well, the size of both the depolarization wave and the repolarization wave are quite similar to the recorded data (see the rightmost panel of Figure 8B).

In theory, the repolarization waves observed in the data and simulations could be used to estimate the action potential duration in the inversion procedure. However, as observed in the lower panels of Figure 8, the repolarization waves are quite smooth, and it is not clear which time points represent different degrees of repolarization. For the V-traces on the other hand (see e.g., Figure 2), it is straightforward to define accurate measures of different degrees of repolarization in the form of APD-values (see Section 2.2.1 and Figure 3). Therefore, we use the optical measurements of V to define the action potential durations and only use the U data for information regarding the depolarization wave.

### 3.4 Estimating drug induced effects on the conduction velocity

One of the advantages of including measurements of the extracellular potential in addition to optical measurements of V and Ca in the inversion procedure is that the extracellular measurements can be used to estimate the conduction velocity of the cell collection. This information could be useful for determining drug effects on the *I*_Na_ current (see Figure 5). The left column of Table 3 reports the conduction velocity computed from measurements of the extracellular potential in a collection of hiPSC-CMs exposed to different doses of the drug Flecainide (measured data, not simulated). We observe that as the drug dose is increased, the conduction velocity is decreased. This could indicate that the drug blocks the *I*_Na_ current, which has also been found in previous studies [5, 7].

**Table 3:** Effect of the drug Flecainide on the conduction velocity computed from the extracellular potential. The left panel reports values based on measurements of hiPSC-CMs and the center and right panels report values computed in bidomain-base model simulations with IC50_CaL_ = 9 *μ*M, IC50_NaL_ = 47 *μ*M, IC50_Kr_ = 1.9 *μ*M, and *h_i_* = 1, for *i*= CaL, NaL, and Kr (see (22)). The drug effect on *I*_Na_ is set to IC50_Na_ = 2.2 *μ*M, *h*_Na_ = 1 in the center column and IC50_Na_ = 0.2 *μ*M, *h*_Na_ = 0.4 in the right column. The parameters used in the simulations are given in Tables 1 and 2, and the default *g*_Na_ value is increased by 45% to match the conduction velocity in the control case.

| Dose | CV data | CV simulation | CV simulation |
| --- | --- | --- | --- |
| | | ($\text{IC50}_{\text{Na}} = 2.2 \mu\text{M}$,<br>$h_{\text{Na}} = 1$) | ($\text{IC50}_{\text{Na}} = 0.2 \mu\text{M}$,<br>$h_{\text{Na}} = 0.4$) |
| 0 $\mu\text{M}$ | 23.7 cm/s | 23.6 cm/s | 23.7 cm/s |
| 1 $\mu\text{M}$ | 17.9 cm/s | 20.0 cm/s | 14.2 cm/s |
| 2.5 $\mu\text{M}$ | 11.7 cm/s | 16.7 cm/s | 12.0 cm/s |
| 10 $\mu\text{M}$ | 9.3 cm/s | 9.2 cm/s | 8.9 cm/s |

In the paper [2], we estimated IC50-values for *I*_CaL_, *I*_NaL_ and *I*_Kr_ based on optical measurements of V and Ca of hiPSC-CMs exposed to Flecainide, but we were not able to estimate the effect on *I*_Na_ due to the low time resolution for the optical measurements. The IC50-values were estimated to IC50_CaL_ = 9 *μ*M, IC50_NaL_ = 47 *μ*M, and IC50_Kr_ = 1.9 *μ*M, where the conductance of each current was scaled according to

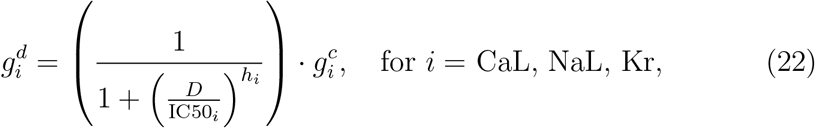

where 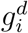 is the conductance of current *i* in presence of the drug dose *D* and 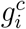 is the conductance in the control case with no drug present. Furthermore, *h*_*i*_ is the so-called Hill coefficient, assumed to be 1 in [2]. Incorporating these IC50-values in addition to some estimated IC50-values for *I*_Na_ in bidomain-base model simulations, we obtain the conduction velocities reported in the center and right columns of Table 3. In the center column we use IC50_Na_ = 2.2 *μ*M and *h*_Na_ = 1, and in the right column we use IC50_Na_ = 0.2 *μ*M and *h*_Na_ = 0.4. Note that since high doses of Flecainide result in increased action potential durations (see e.g., [2]), we have used a pacing frequency of 0.5 Hz instead of 1 Hz when updating the initial conditions for these simulations. In addition, to match the conduction velocity in the control case, the default value of *g*_Na_ value of Table 2 is increased by 45%. We observe that the bidomain-base model simulations are able to roughly reproduce the drug induced reduction in conduction velocity observed in the measurements, indicating that a comparison of measured and simulated conduction velocities could help identify drug effects on *I*_Na_.

The data and model solutions are further compared in Figure 9. Here, we show the U solutions and the measured U data in the 64 electrodes for the control case and for the 10 *μ*M dose case at some different points in time. In the simulations, we use IC50_Na_ = 0.2 *μ*M and *h*_Na_ = 0.4, and we have adjusted the stimulation location to correspond to the propagation direction observed in the measured data. In the control case, the wave moves in the *y*-direction from the lower part of the domain to the upper part, and in the drug case, the wave moves from the upper left corner to the lower right corner of the domain. In both the data and the simulations, we observe that the extracellular depolarization wave moves more slowly across the domain for 10 *μ*M Flecainide than in the control case, consistent with the reduced conduction velocities observed in Table 3.

**Figure 9:**
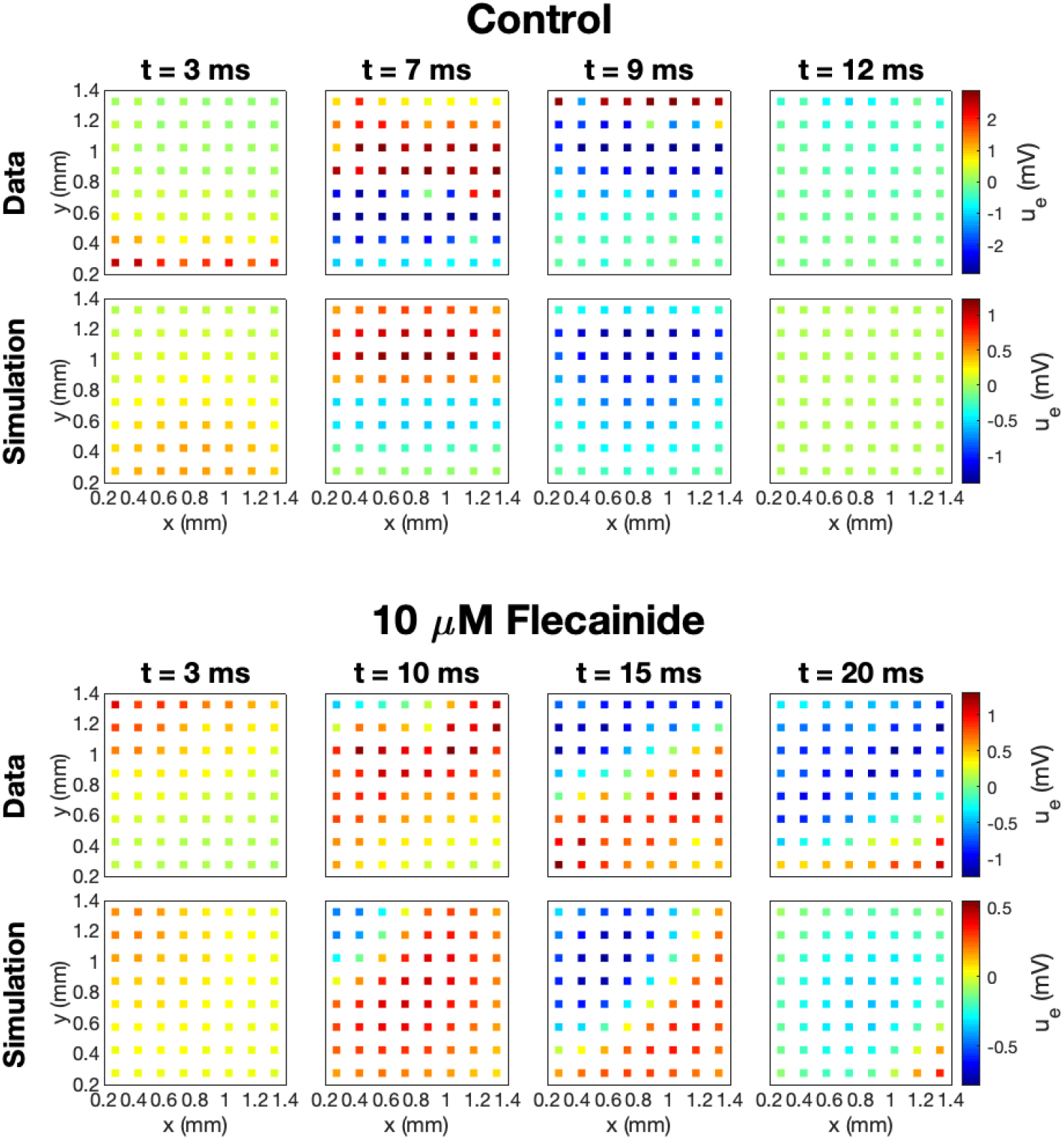
Measured and simulated U data in 64 electrodes for the control case and for 10 *μ*M Flecainide, modeled using IC50_CaL_ = 9 *μ*M, IC50_NaL_ = 47 *μ*M, IC50_Kr_ = 1.9 *μ*M, IC50_Na_ = 0.2 *μ*M, *h_i_* = 1 for *i* = CaL, NaL, and Kr, and *h*_Na_ = 0.4. The parameters used in the simulations are given in Tables 1 and 2, and the default *g*_Na_ value is increased by 45%. The stimulation location is adjusted to correspond to the propagation direction in the data.

### 3.5 Inversion of simulated drugs

In order to investigate how well the inversion procedure outlined above is able to identify the effect of drugs on *I*_Kr_, *I*_CaL_ and *I*_Na_, we generate simulated data for twelve drugs whose effect on *I*_Kr_, *I*_CaL_ and *I*_Na_ was investigated in [7]. The block percentages used to generate the data are based on the block percentages for drug concentrations corresponding to three times the free plasma *C*_max_ reported in [7]. However, large block percentages for *I*_Kr_ have been reduced in order to obtain reasonable V and Ca traces. In the inversions, we use two extracellular calcium concentrations, as explained in Section 2.3. Furthermore, we save U data and information about the Ca peak time extracted from the first 50 ms of the bidomain simulations for all the drugs, except for the drug Diltiazem. For Diltiazem we had to increase the bidomain simulation time to 100 ms in order to ensure that some of the grid points in the domain had reached their peak calcium concentration to compute the Ca peak time (see Section 2.2.1).

In Figure 10, we compare the block percentages estimated by the inversion procedure to those used to generate the data for the twelve drugs. We observe that for all the considered drugs, the inversion procedure is able to identify the block of the three currents quite accurately.

**Figure 10:**
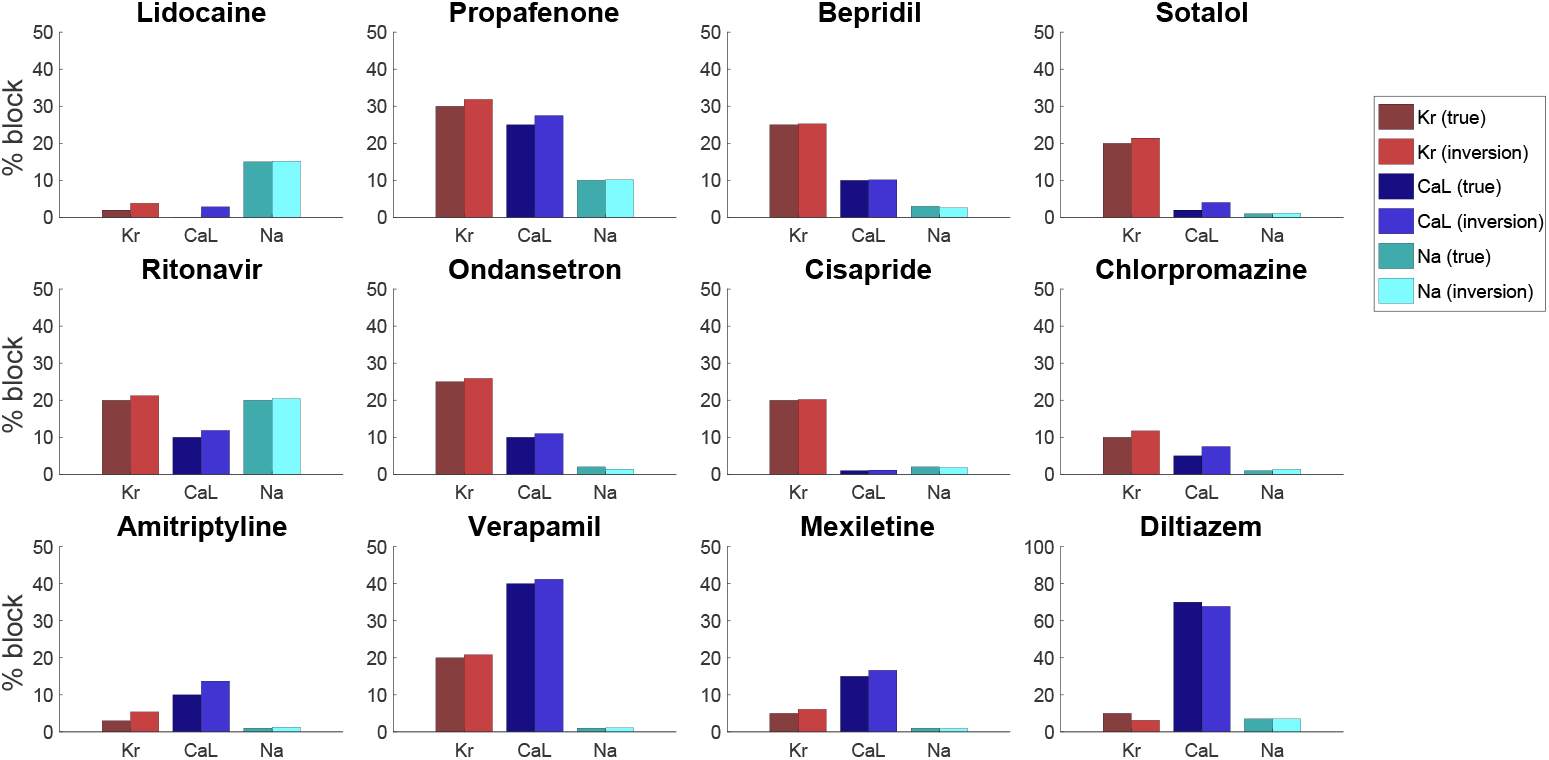
Result of the inversion procedure applied to simulated data of twelve drugs, based on the block percentages corresponding to three times the free plasma *C*_max_ reported in [7].

### 3.6 Effect of noise

In Figure 10, we considered how well the inversion procedure was able to identify drugs based on data generated by bidoman-base model simulations, and we observed that the inversion procedure was able to identify the correct channel blocking quite accurately. However, when the V, Ca and U data are recorded from real measurements from microphysiological systems, the data will include some noise. In order to investigate how well the inversion procedure is able to identify the effect of drugs from data including noise, we include 5% noise in the V data, 3% noise in the Ca data and 1% noise in the U data and repeat the inversions shown in Figure 10. The noise is added by drawing a random number between −*p · A* and *p* · *A* for each point in time, where *A* is the difference between the maximum and minimum value of the considered data and *p* is the noise percentage. This random number is then added to the V, Ca and U traces.

The result of the inversions with noise included in the data is given in Figure 11. We observe that the inversion procedure is able to estimate the block percentage quite accurately in most cases, but that the accuracy is reduced compared to the case with no noise in Figure 10.

**Figure 11:**
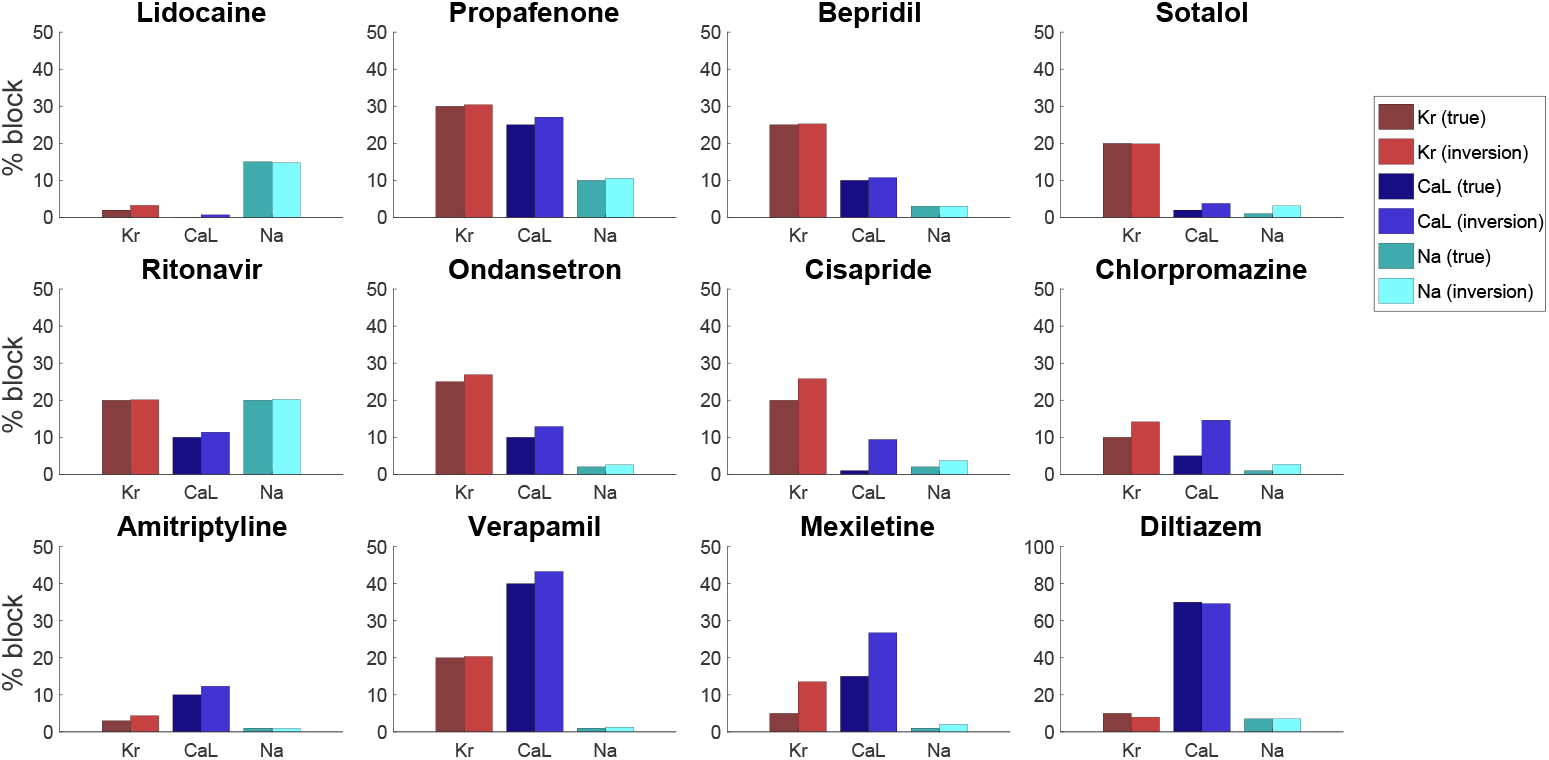
Result of the inversion procedure applied to simulated data of the twelve drugs of Figure 10. We have here included 5% noise in the V data, 3% noise in the Ca data and 1% noise in the U data.

## 4 Discussion

### 4.1 Sensitivity of parameters is necessary for identification

A generic problem in mathematical models of physiology is to determine the parameters included in the model. For models of the action potential of excitable cells, there is a large number of parameters that needs to be determined in order to use the model; see e.g., [33, 34, 35, 36]. If a model is insensitive to changes in a specific parameter, that parameter is impossible to determine by comparing experimental results and results of numerical simulations. Contrarily, if the model is sensitive to a parameter, numerical experiments can be used to match the model to the data by fine-tuning the parameter. In Figure 4 above, we saw that the membrane potential is clearly sensitive to changes in the *I*_Kr_ and *I*_CaL_ currents. The upstroke is also sensitive to changes in the *I*_Na_ current, but since the upstroke is very fast, the sensitivity is difficult to see in the plot of the entire AP. Since optical measurements have a relatively coarse time resolution, we have been unable to identify the strength of the sodium current based solely on voltage and calcium traces. However, we note from Figure 4 that the extracellular potential is indeed sensitive to changes in the sodium current. This sensitivity also carries over to sensitivity of the conduction velocity (CV) with respect to changes in the sodium current, and this sensitivity is utilized in the cost function; see (15).

### 4.2 The conduction velocity is governed by the sodium current

The conduction of the electrochemical signal through the cardiac muscle is essential for the functioning of the heart. Surprisingly, the conduction process is still not completely understood; see, e.g., [37] for a review of the development of the understanding of cardiac conduction. Globally (mm scale), cardiac conduction is usually modeled using homogenized models like the bidomain or monodomain models (see e.g., [17, 22, 23]), but locally (*μ*m scale) more detailed models are used; see e.g., [38, 39, 40]. The microphysiological system is somewhere in between these scales (the size is about 1.2 mm × 1.2 mm) and we would have liked to use the detailed EMI model introduced for cardiac conduction in [41]. However, the EMI approach is challenging from a computational point of view and we have therefore used the much simpler bidomain model to consider how the CV depends on changing the different ion currents.

In Figure 5 we observe that the CV is very sensitive to changes of the *I*_Na_ current, but almost insensitive to changes in the *I*_Kr_ and *I*_CaL_ currents. This means that we can use the measurements of U to identify the *I*_Na_ current first and then only look for the *I*_Kr_ and *I*_CaL_ currents based on the V- and Ca-traces.

### 4.3 Improving visibility of parameters by changing the extracellular concentrations

In using experimental data to determine parameters of a computational model, it is desirable to use protocols designed to highlight the effect of different parameters. For instance, it is argued in [42] that random stimulation protocols can improve visibility of the parameters, and this is confirmed in [36]. Another experimental parameter that can be changed is the ionic concentrations of the extracellular environment. In Figure 6 and Figure 7 we show that using two different extracellular calcium concentration can improve identifiability of the model parameters. One particular important effect of this is that multiple extracellular calcium concentration can aid in distinguishing changes of the *I*_Kr_ and *I*_CaL_ currents. Since blocking *I*_Kr_ has more or less the same effect as increasing *I*_CaL_, it is difficult to distinguish these effects in measurements of the AP. But it is apparent from Figure 7 that the level of blocking changes with the extracellular calcium concentration, and this can be used to distinguish between different blocking combinations for these two currents.

### 4.4 Inversion of simulated drugs

Simulated data is often used as a proxy for real data when the real data are cumbersome to obtain. Here, we have used real data for U both for the case of no drug and when various doses of Flecainide have been applied. Furthermore, the base model used to represent the membrane dynamics has been parameterized using real data; see [2, 1]. However, in order to study inversion for the range of data provided in the CIPA report [7], we have used simulated data. One advantage of this is that we can get data to any desired accuracy and that we can control the level of noise inevitably introduced in the measurements. The disadvantage is clearly the reduced realism of the data and it is a priority of future work to use combined V, Ca, and U data to do inversion of both well characterized and novel drugs with hitherto unknown properties.

For simulated data, we notice that the inversion procedure provides quite accurate estimates. Systematically, we observe a tendency to overestimate the block of both the *I*_Kr_ and *I*_CaL_ currents. As alluded to above, it is notoriously difficult to distinguish reduced *I*_Kr_ from increased *I*_CaL_ and vice versa. This can be improved (Figure 6 and Figure 7), by using several values of the extracellular calcium concentration, but the problem is not completely removed.

### 4.5 Repolarization waves in bidomain simulations

As mentioned above, the bidoman model has been used by many authors (see e.g., [18, 16, 19, 20, 14]) to simulate the electrophysiology of collections of hiPSC-CMs. One problem pointed out by several authors is the lack of a repolarization wave in the simulated results although the repolarization wave is clearly present in measurements of the extracellular potential; see, e.g., Figure 3 and Figure 4 of [18]. This feature of the bidomain solution is repaired by introducing heterogeneities in the tissue which lead to a repolarization wave. In Figure 8A, we show that there is indeed a repolarization wave present in the extracellular data obtained from collections of hiPSC-CMs. The repolarization wave is also present in our bidomain simulations, but the strength of the wave depends critically on the intracellular conductivity, *σ*_*i*_. The repolarization wave becomes stronger as *σ*_*i*_ is reduced. Since the intracellular conductivity represents the geometrical average of the intercellular conductivity (regulated by gap junctions) and the cytosolic conductivity, it is reasonable to use a reduced value of *σ*_*i*_ for immature cells, since the gap junctions are most likely less developed in collections of hiPSC-CMs than for collections of adult cardiomyocytes. In [18], it is argued that it is reasonable to introduce heterogeneities in the tissue and we agree. However, our results indicate that it is not necessary in order to see a repolarization wave in the simulation results.

## 5 Conclusion

We have shown that by using data traces of the membrane potential, the intracellular calcium concentration and the extracellular potential, we can estimate the major sodium, calcium and potassium currents in the base model. It remains to enable concurrent observation of all three modalities, but when such data become available, the methodology described in the present report may be used to invert the data and thus obtain channel densities and estimate drug effects on the channels.

## Notes

### Competing Interest Statement

KHJ, SW, KH, and AT have financial relationships with Organos Inc, and the company may benefit from commercialization of the results of this research.

